# The placental protein NRK promotes cell death through its plasma membrane-localizing CNH domain

**DOI:** 10.1101/2023.12.27.573405

**Authors:** Beni Lestari, Kohei Soda, Kei Moritsugu, Akinori Kidera, Yusuke Suenaga, Yoshitaka Hippo, Edy Meiyanto, Masayuki Komada, Mae Sri Hartati Wahyuningsih, Toshiaki Fukushima

## Abstract

Placental development is regulated by the balance between cell proliferation and death. The placental protein NRK (NIK-related kinase) plays a role in preventing excessive placenta growth. We previously demonstrated that NRK underwent rapid molecular evolution in the ancestor of placental mammals and acquired the functional regions, including the phospholipid-binding citron homology (CNH) domain, by which NRK inhibits cell proliferation. NRK is also potentially responsible for cell death; caspases cleave NRK during apoptosis, releasing the *C*-terminal fragment that promotes cell death. Here, we explored the molecular mechanisms underlying the cell death-promoting effects of NRK. Our experimental data using HeLa, placenta trophoblast BeWo (human), and Rcho-1 (rat) cells indicated that the CNH domain of NRK was required and sufficient to promote cell death. *In vitro* and *in silico* studies showed the NRK CNH domain bound to phospholipids via its polybasic clusters and remains at the plasma membrane (PM) during apoptosis. Evolutional analyses indicated that these clusters formed in the ancestor of placental mammals. Mutations in these clusters (CNH-18A) hindered the cell death-promoting activity of the CNH domain. We concluded that NRK promotes cell death through its plasma membrane-localizing CNH domain and suggested its active role in PM-associated events during cell death.

## Introduction

The placenta is the first and largest fetal organ to develop after conception and is essential for the success of eutherian reproduction (1–3). Placental development is complex and regulated by many factors to balance cell proliferation, differentiation, and death (4–6). Disorders related to the placenta impact approximately one-third of human pregnancies, leading to significant maternal and fetal mortality and morbidity (7, 8). Despite its critical role, the intricate mechanisms governing placental development and functional regulation remain largely unknown.

Apoptosis is a process important for normal development and tissue homeostasis (9, 10). Morphological hallmarks of apoptosis include cell shrinkage and rounding, membrane blebbing, chromatin condensation, DNA fragmentation, and formation of apoptotic bodies (11). The incidence of apoptosis is highest at the last stage of placental development in humans (12–15). In rats, the placental spongiotrophoblast layer, the middle junctional area with endocrine and nutrient storage functions, exhibits a higher level of apoptosis than other layers at the late stage of pregnancy (16). These observations suggest that apoptosis is important in placental remodeling and maturation during late pregnancy.

NIK-related kinase (NRK), also called NIK-like embryo-specific kinase (NESK), is a serine/threonine kinase encoded by the *Nrk* gene on the X chromosome (17). The *Nrk* gene is located in the X chromosome region with the highest accelerated evolution in the ancestor of placental mammals (18). In mice, NRK is expressed highly in the placental spongiotrophoblasts and moderately in the trophoblast giant cells and syncytiotrophoblasts-2 in the labyrinth layer during the mid to late stage of pregnancy (19, 20). In the human placenta, NRK is expressed in cytotrophoblasts and syncytiotrophoblasts in villi (21), which share similar functions to the mouse spongiotrophoblast and labyrinth layers, respectively (1, 22, 23). *Nrk* KO mice showed placental hyperplasia with a much enlarged spongiotrophoblast layer and moderately enlarged labyrinth layer (19, 24), demonstrating the role of NRK in preventing excessive placenta growth.

NRK is a member of the Germinal Center Kinase (GCK) family subgroup IV with a three-domain structure: an *N*-terminal kinase domain, a middle disordered region, and a *C*-terminal citron homology (CNH) domain (25, 26). We previously demonstrated that NRK underwent rapid molecular evolution in the ancestor of placental mammals and acquired the functional regions, including the CK2 inhibitory region (CIR) in the middle region and the phospholipid-binding CNH domain (20). Through the CIR and CNH domain, NRK is located at the plasma membrane (PM) and inhibits proliferative signal via the CK2-PTEN-AKT pathway (20), suggesting its role in suppressing placental cell over-proliferation.

On the other hand, NRK is a putative cell death regulator. In HeLa cells exogenously expressing mouse NRK (mNRK), caspases cleave mNRK at Asp-868 and −1091 residues upon the induction of apoptosis by TNF-α and CHX. The *C*-terminal fragment containing the CNH domain has an activity that makes cells more susceptible to cell death (27). These imply that NRK may be involved in apoptosis to support proper placenta size. This study explored the molecular mechanisms by which NRK promotes cell death and the underlying molecular evolution.

## Results

### The responsible region for the cell death-promoting activity of mNRK

To identify the region of mNRK responsible for its cell death-promoting activity, a series of deletion mutants of mNRK (Fig 1A) were transiently expressed in HeLa cells and treated with the apoptotic inducers, TNF-α and CHX for 6 h. In the negative control sample (mock), approximately 85% of cells were viable. The expression of full-length mNRK, the amino acid (aa) 1092-1455 fragment, and the CNH domain significantly decreased cell viability, with a reduction of to 60-70%. The mutant lacking the CNH domain (ΔCNH) failed to decrease cell viability (Fig 1B). We also generated HeLa cells stably expressing the CNH domain and subjected the cells to apoptotic inducers for various durations. mNRK CNH domain-expressing cells exhibited a significant decrease in cell survival compared to the parental cells (Fig. 1C). These data indicated that the CNH domain is required and sufficient for the cell death-promoting activity of mNRK.

**Figure 1.**
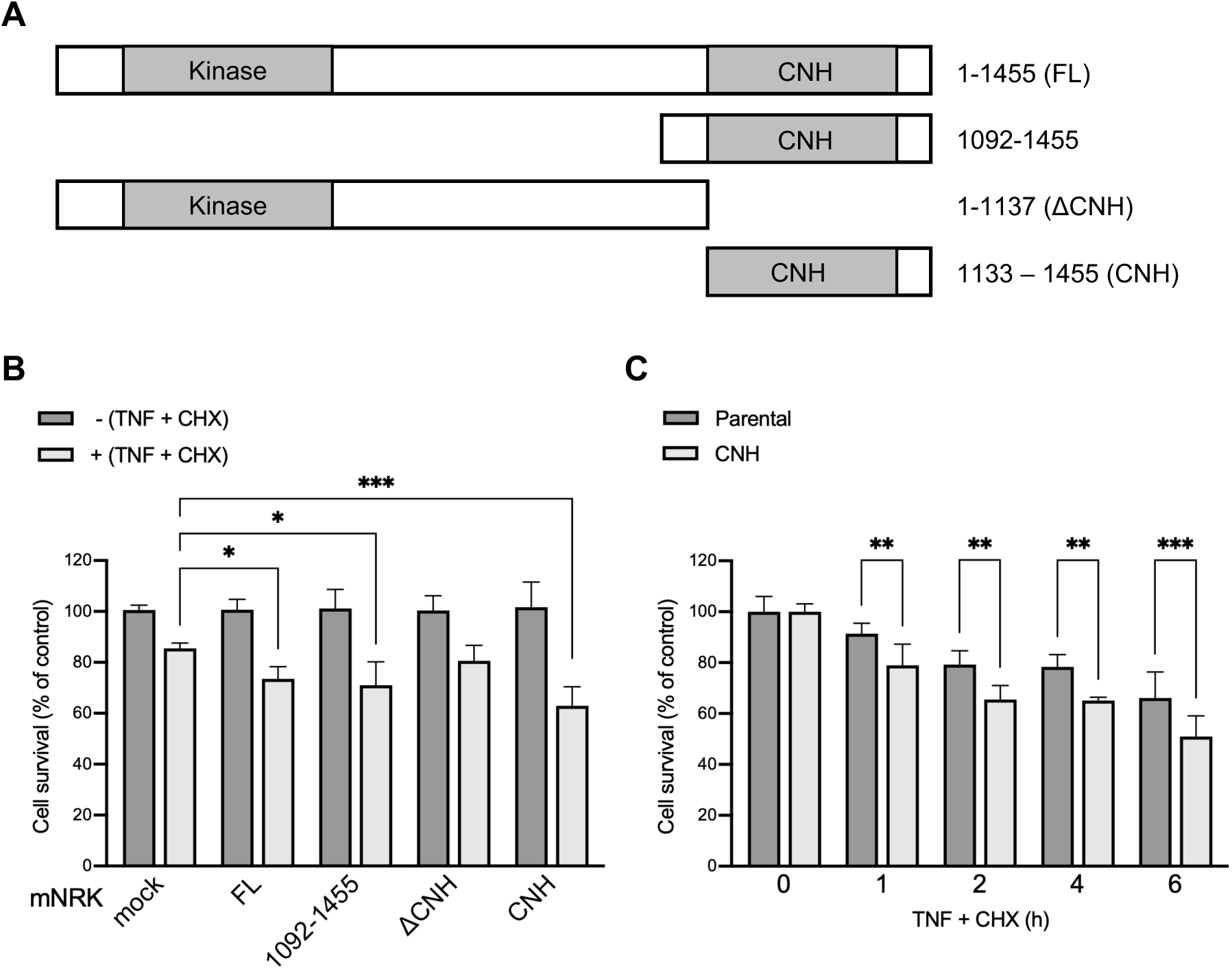
The cell death-promoting activity of mNRK CNH domain. *A-C*, HeLa cells were transfected with plasmids encoding indicated proteins (A), or cells stably expressing the CNH domain via lentivirus were generated (C). The cells were stimulated with apoptotic inducers (10 ng/mL TNF-α and 10 µg/mL CHX) for 6 h (B) or indicated durations (C) and then subjected to WST-8 assay. Experiments were performed in triplicate or quadruplicate. The graph shows the mean ± standard deviation (SD) of all experiments. Statistical significance was determined through one-way analysis of variance (ANOVA) followed by Dunnett’s multiple comparisons test: **** p ≤ 0.0001; *** p ≤ 0.001; ** p ≤ 0.01; * p ≤ 0.05.

### The localization of the mNRK CNH domain in cells undergoing apoptosis

The mNRK CNH domain exhibits PM localization under normal cell culture conditions (20). We assessed its localization during the progression of apoptosis. Cells were treated by TNF-α and CHX, and differential interference contrast (DIC) micrographs were used to distinguish the different stages of apoptosis. The CNH domain retained its PM localization in pre-apoptotic cells with shrinking morphology. Furthermore, it accumulated at the membrane blebbing sites in both early- and late-apoptotic cells, characterized by a few and many blebs, respectively (Fig. 2A). These findings suggest that apoptotic induction does not affect the localization of the CNH domain, suggesting an active role of this domain in PM-associated events during cell death.

**Figure 2.**
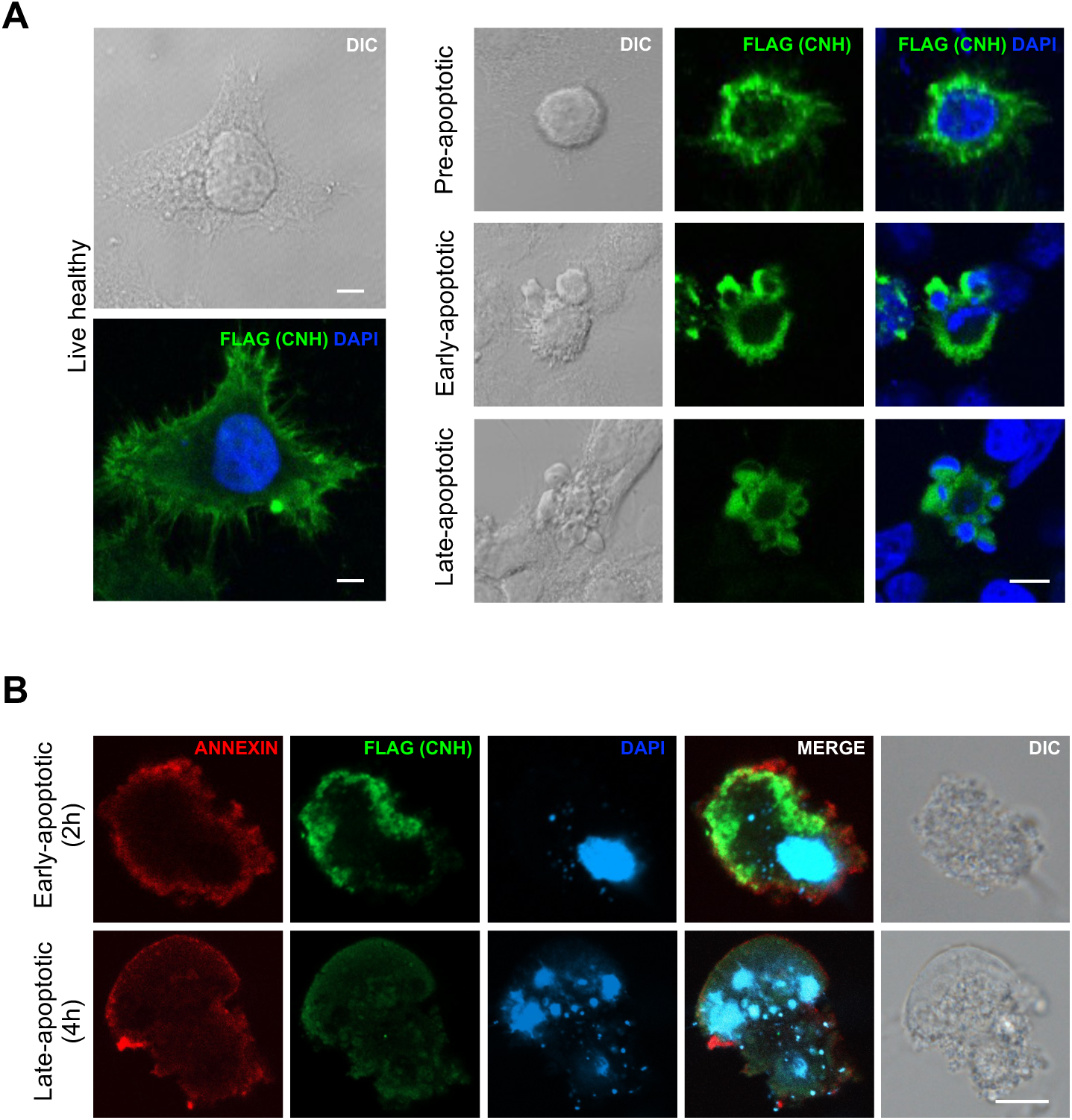
Localization of the mNRK CNH domain during apoptosis. *A*, HeLa cells were transfected with plasmids encoding FLAG-tagged mNRK CNH domain and treated with apoptotic inducers (10 ng/mL TNF-α and 10 µg/mL CHX) for 0, 2, and 4 h. Cells were subjected to fixation, permeabilization, and immunostaining. Nuclei were stained with DAPI. *B*, After apoptotic induction for 2 and 4 h, cells were incubated with Annexin V fluorescent conjugates and subjected to fixation and immunostaining for the CNH domain. Scale bar, 10 μm.

Phosphatidylserine (PS) is typically confined to the inner leaflet of the cell membrane in living cells, and its exposure on the outer leaflet occurs during apoptosis (28). The mNRK CNH domain can bind to membrane phospholipids, specifically PS (20). We stained PS of cells with Annexin V fluorescent conjugates and then performed immunostaining for the CNH domain. In this experiment, cells were not permeabilized, and the FLAG-positive cells should have lost membrane integrity due to the ongoing secondary necrosis (29). The result showed that the mNRK CNH domain is partly localized in proximity to PS (Fig. 2B), further supporting PM localization of the mNRK CNH domain in dead cells.

### The molecular basis for the PM localization of the mNRK CNH domain

To determine sites of the CNH domain responsible for PM localization, we compared the localization of the GFP-tagged full-length CNH domain (GFP-CNH FL, aa 1133-1455) and truncated mutants, GFP-CNH A (aa 1133-1331) and GFP-CNH B (aa 1332-1455), using immunostaining. The truncation site was in the loop following the fourth β-propeller, which was determined based on the structural model of the CNH domain described below. The results revealed that both truncated mutants displayed a weakened PM localization compared to GFP-CNH FL (Fig. 3A), indicating that both regions are involved in membrane localization.

**Figure 3.**
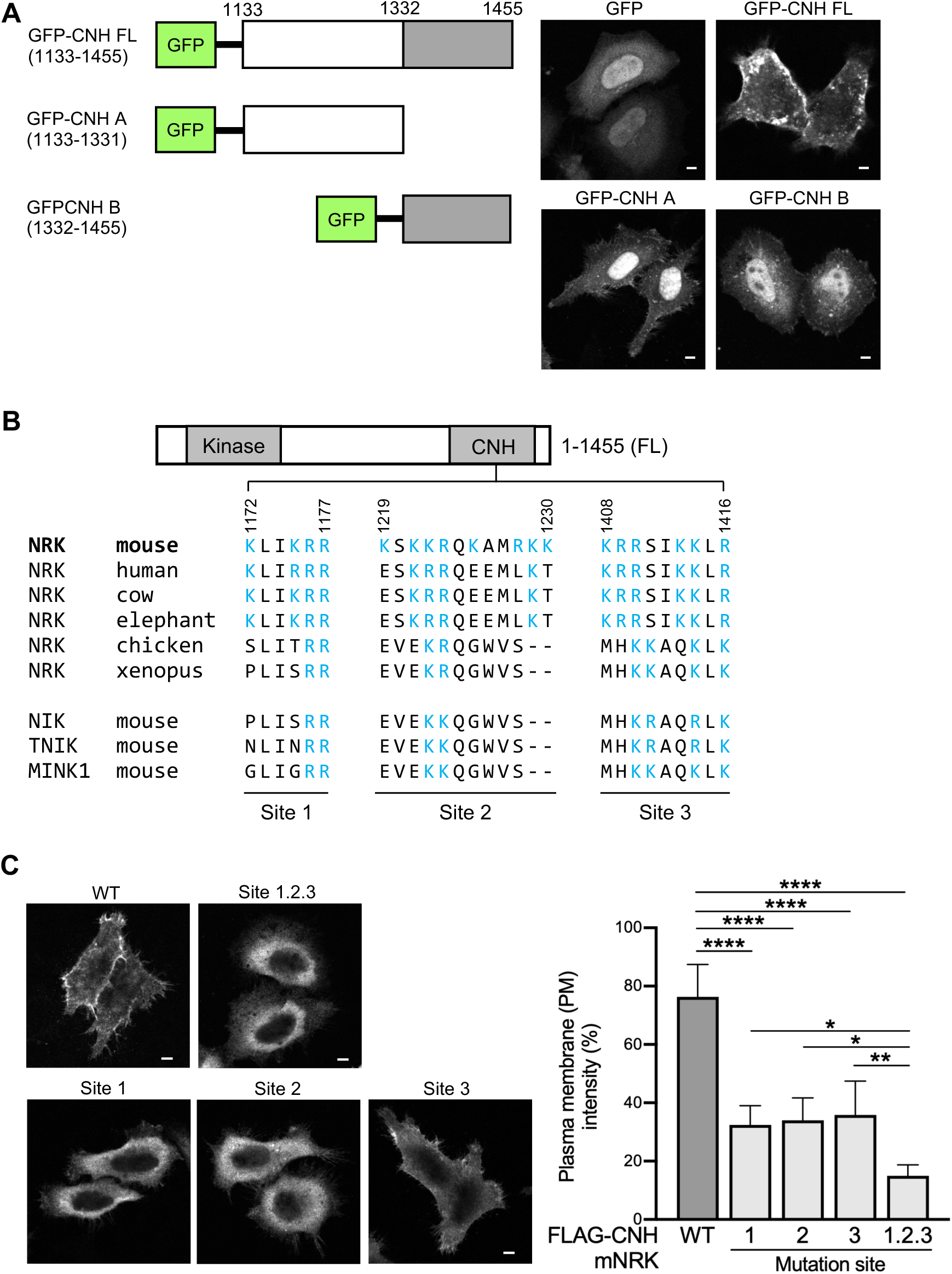
Responsible sites of the mNRK CNH domain for plasma membrane localization. *A,* Schematic structure of GFP-tagged truncated forms of the CNH domain (left panel). Fluorescence microscopy analysis of HeLa cells expressing the indicated proteins (right panel). *B,* Sequence alignment of polybasic clusters (sites 1-3) within the NRK CNH domain across different species and other GCK IV kinases. The blue color indicates the presence of basic or positively charged amino acids, specifically lysine (K) and arginine (R). *C,* Basic amino acids in sites 1-3 within the FLAG-tagged CNH domain were mutated to alanine. HeLa cells expressing proteins mutated at the indicated sites were subjected to immunostaining with an anti-FLAG antibody. PM fluorescence intensity was measured using NIS-Elements AR analysis. The graph represents the mean ± standard deviation (SD) of five cells. Statistical significance was determined through one-way analysis of variance (ANOVA) followed by Dunnett’s multiple comparisons test: **** p ≤ 0.0001; ** p ≤ 0.01; * p ≤ 0.05.

Subsequent sequence analysis of the mNRK CNH domain revealed three sites rich in positively charged amino acids, conserved only in eutherian NRK (site 1, aa 1172-1177; site 2, aa 1219-1230; site 3, aa 1408-1416) (Fig. 3B). To determine whether these polybasic clusters are involved in PM localization of the mNRK CNH domain, we made the mutants in which all basic residues in each site were substituted by alanine. Immunofluorescence analysis showed that mutation in each site reduced PM localization up to 50% to that of WT mNRK CNH. The mutant in which all eighteen basic residues in sites 1-3 were substituted showed the lowest PM intensity (Fig. 3C), indicating that all polybasic clusters are essential for PM localization.

We further investigated the effects of mutations at the polybasic clusters on the *in vitro* binding of the mNRK CNH domain to PS-coated beads. FLAG-tagged CNH domains with or without mutations in sites 1-3 were produced in HEK293T cells and purified using FLAG-antibody. The PS-coated beads binding assay showed that the mutations of each site did not affect PS-binding, and the mutation of all sites significantly reduced it (Fig. 4). These results suggest that the basic residues at sites 1-3 are essential for PS binding. Single-cluster mutations partially reduced PM localization (Fig. 3C) but did not affect PS-binding (Fig. 4), likely because PS enriched on beads can adsorb the CNH domain more effectively than plasma membrane lipids. Taking them together with the data above, we concluded that the mNRK CNH domain localizes at the PM via its polybasic clusters by binding to phospholipid (mainly PS).

**Figure 4.**
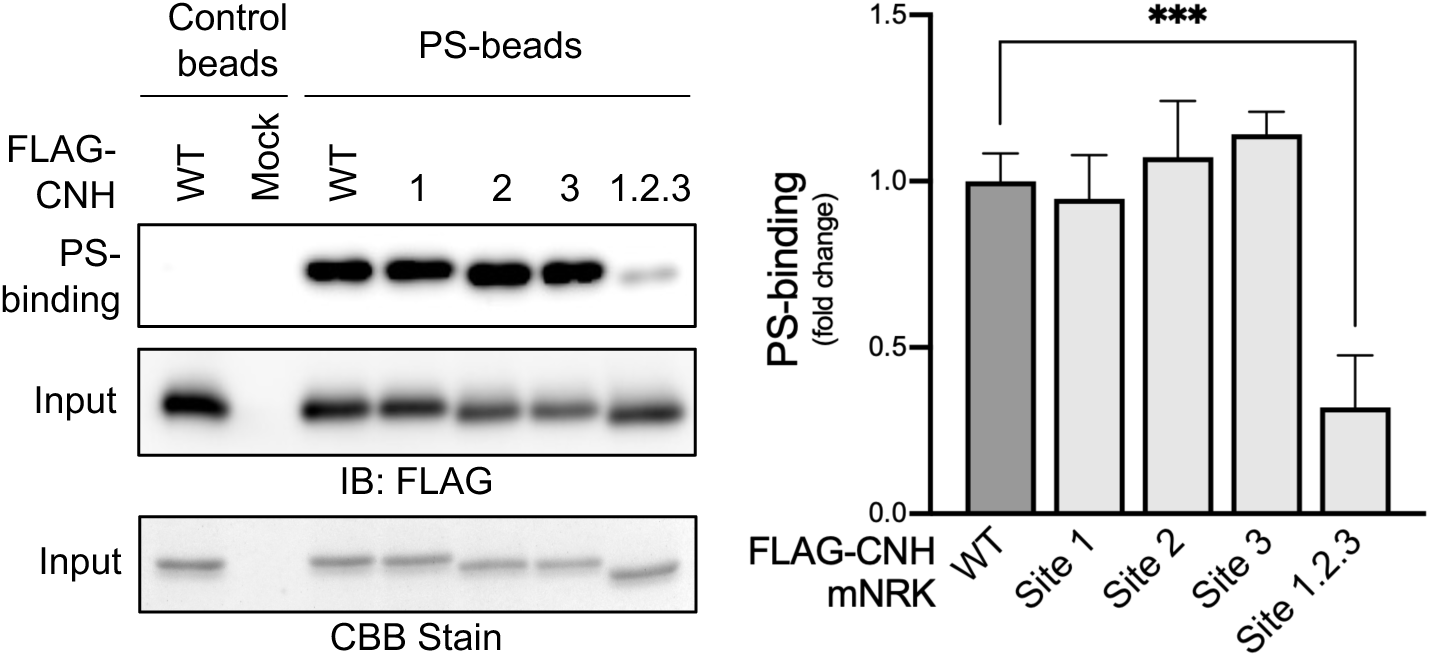
Phosphatidylserine-binding ability of the mNRK CNH domain and the involvement of the polybasic clusters. HEK293T cells were transfected with plasmids encoding FLAG-tagged mNRK CNH domain (wild-type and mutants of sites 1-3). The CNH domains were purified by immunoprecipitation with anti-FLAG antibody and elution with FLAG peptide. Protein purification was confirmed by Coomassie Brilliant Blue (CBB) staining. Purified proteins were subjected to phosphatidylserine (PS)-binding assay and immunoblotting with anti-FLAG antibody. The densitometric analyses were performed. The graph represents mean values ± standard deviation (SD) from three independent experiments. Statistical significance was determined through one-way analysis of variance (ANOVA) followed by Dunnett’s multiple comparisons test: *** p ≤ 0.001.

### *In-silico* structural modeling of the CNH domain of mNRK and chicken NRK (cNRK) and their interactions with lipid bilayer

We constructed an *in-silico* structural model of the mNRK CNH domain. Since the chicken NRK (cNRK) CNH domain cannot bind to phospholipids *in vitro* (20), we also constructed a model of the cNRK CNH domain for comparison. These models show a β-propeller structure similar to the Rom2 CNH domain (30), with substantial stability (supplementary figure 1). Sites 1-3 of the polybasic clusters in the mNRK CNH domain (colored in blue, light, and cyan) seem to be on the same surface extending perpendicular to the central tunnel. On the opposite surface, the mNRK CNH domain has a structure containing two α-helixes (colored in yellow) not seen in the cNRK CNH domain. It is encoded by mouse *Nrk* exon 25, an exon found only in eutherian *Nrk* genes (20). The surface containing sites 1-3 of the mNRK CNH domain had a larger area of negative charge than the corresponding surface of the cNRK CNH domain (Fig. 5A, B).

**Figure 5.**
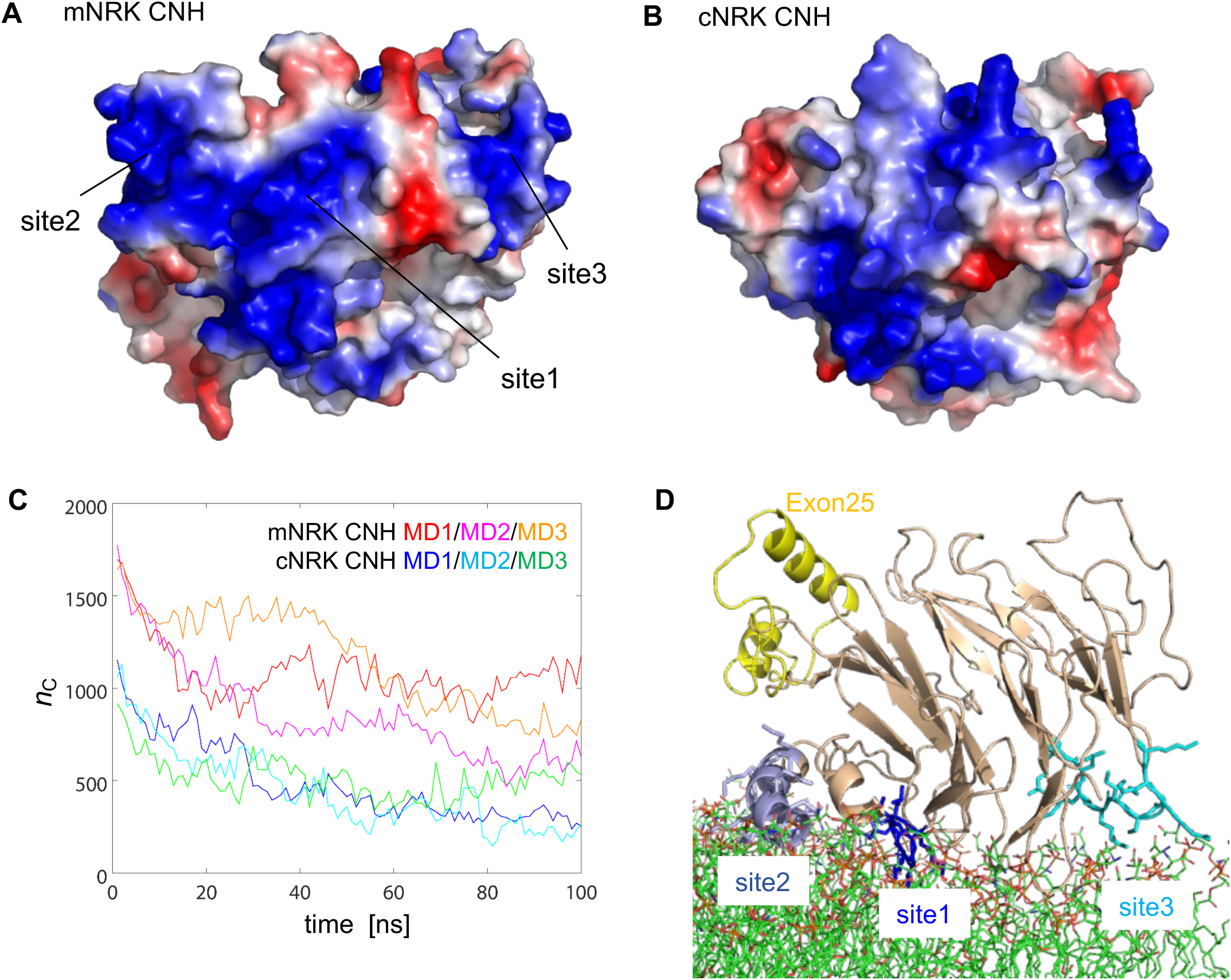
*In silico* structural analyses of the interactions between the CNH domains and lipid bilayer. *A,B,* The electrostatic potential of the surface containing sites 1-3 of the mNRK CNH domain (A) and the corresponding surface of the cNRK CNH domain (B) are shown. *C,* Using the structural models of CNH domains and palmitoyl-oleoyl phosphatidylserine (POPS) lipid bilayer, the initial structures interacting with lipid bilayer were constructed, and these MD simulations were performed three times (MD1-3). The number of atom contacts between the CNH domains and lipid bilayer (with the interatomic distance ≤ 5 Å) is shown. *D,* Representative structure of the mNRK CNH domain interacting with the phospholipid is taken from MD1.

We performed *in silico* structural analyses of the interaction between the CNH domains and a palmitoyl-oleoyl phosphatidylserine (POPS) lipid bilayer. Compared to the cNRK CNH domain, the mNRK CNH domain has more atom contacts with the lipid bilayer in the initially predicted structures, and two out of three MD simulations maintained stable contacts in which sites 1-3 interact with the lipid bilayer (Fig. 5C, D, and supplementary figure 2A). In contrast, the cNRK CNH domain lost most atom contacts during simulation (Fig. 5C and supplementary figure 2B). These results suggest that the polybasic clusters in the mNRK CNH domain function as contact sites with phospholipids in the plasma membrane.

### The requirement of the polybasic clusters for the cell death-promoting activity of the mNRK CNH domain

Next, we investigated the role of the phospholipid-binding ability of the mNRK CNH domain in cell death progression. We generated HeLa cells stably expressing the CNH domain mutant in which all eighteen basic residues in three polybasic clusters were substituted by alanine (hereafter referred to as CNH-18A). Analysis of cell death and survival revealed that, unlike the intact CNH domain, CNH-18A expression did not affect cell death progression (Fig. 6A and B). We obtained similar results in the experiments using human and rat trophoblast cell lines BeWo (31, 32) and Rcho-1 (33, 34) (Fig. 6C and D). These results indicate the critical role of the phospholipid-binding ability of the CNH domain in promoting cell death in various types of cells, including trophoblasts.

**Figure 6.**
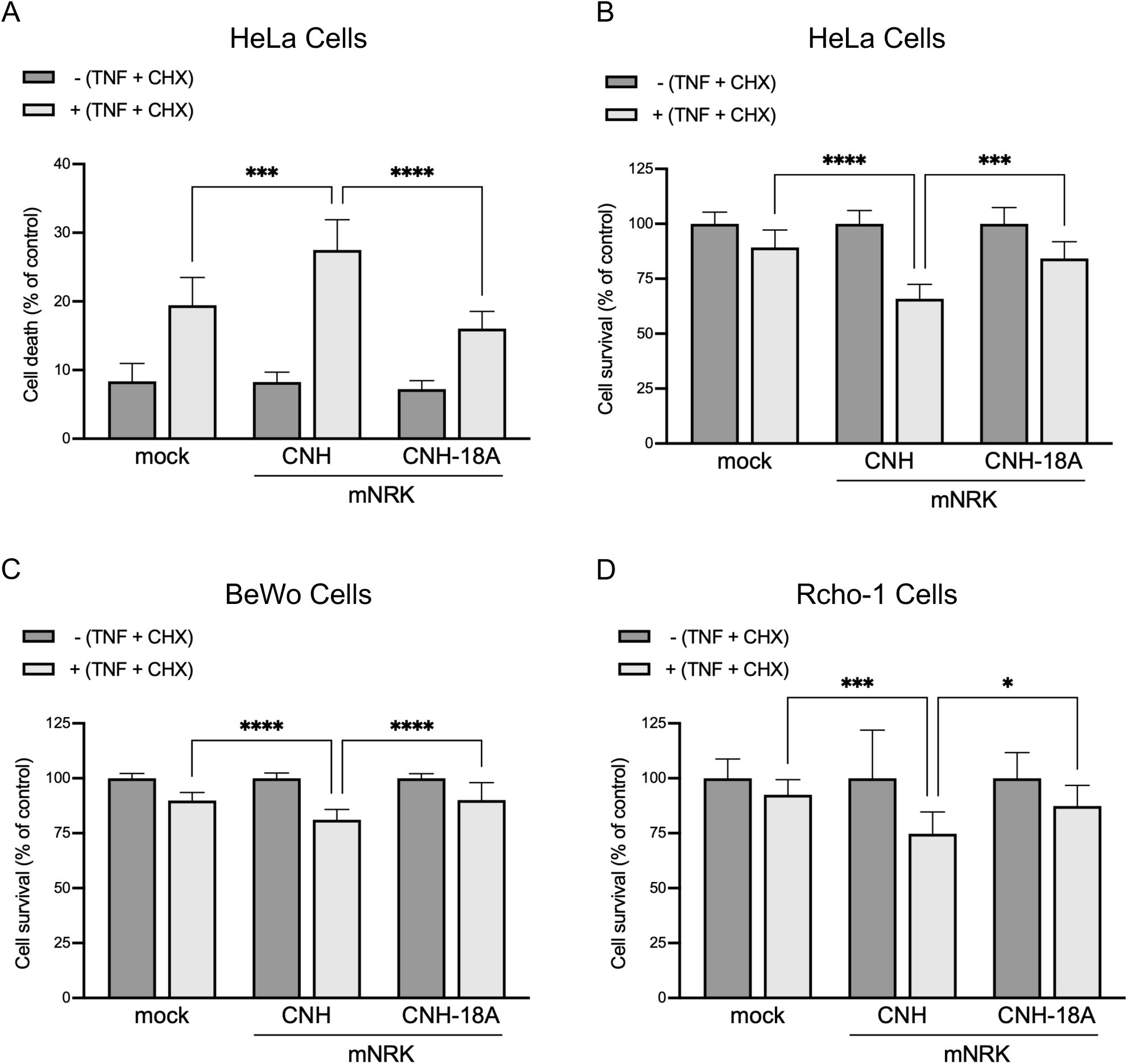
The role of the polybasic clusters in the cell death-promoting activity of the mNRK CNH domain. *A, B,* HeLa cells stably expressing the mNRK CNH domain and CNH-18A were stimulated with apoptotic inducers (10 ng/mL TNF-α and 10 µg/mL CHX) for 6 h and then subjected to WST-8 assay (A) and SYTOX Green staining (B). *C, D,* BeWo (C) and Rcho-1 (D) cells transiently expressing the mNRK CNH domain and CNH-18A were stimulated with apoptotic inducers (10 ng/mL TNF-α and 10 µg/mL CHX) for 6 h and then subjected to WST-8 assay. Statistical significance was determined through one-way analysis of variance (ANOVA) followed by Dunnett’s multiple comparisons test: **** p ≤ 0.0001; *** p ≤ 0.001; ** p ≤ 0.01; * p ≤ 0.05.

The expression of the mNRK CNH domain did not affect caspase-3, PARP1 cleavage, and JNK phosphorylation upon apoptosis induction (supplementary figure 3A), suggesting that the CNH domain may not act upstream of the caspase and JNK pathway. To explore effector protein(s) of the CNH domain, we performed co-immunoprecipitation and mass spectrometry analysis and identified 14 CNH domain-interacting proteins, including four PM-localizing proteins (supplementary figure 3B, supplementary table 1).

## Discussion

We demonstrated that the NRK CNH domain underwent molecular evolution to acquire polybasic clusters that facilitate binding to phospholipids at the PM. Upon apoptosis induction, caspases cleave mNRK to generate the C-terminal fragment containing the CNH domain. We found that the CNH domain remains at the PM during apoptosis progression, interacts with some proteins, and promotes cell death. Although the mechanisms by which the CNH domain promotes cell death remain unclear, we showed that the phospholipid-binding ability was crucial.

### Molecular evolution and structure of the NRK CNH domain

The *C-*terminal CNH domain is conserved among all four GCK IV family members (NIK, TNIK, MINK1, and NRK). Although the CNH domains of NIK, TNIK, and MINK1 are reported to interact with Rap2A, a PM-localizing small GTPase (35–38), they rarely localize at the PM (39). The mNRK CNH domain does not interact with Rap2A (27), but it displays clear localization at the PM. This ability appears exclusive to the CNH domain of eutherian NRK because cNRK does not localize at the PM (20). This study identified three polybasic clusters within the mNRK CNH domain responsible for interacting with phospholipids and PM localization. Other GCK IV family members and non-eutherian NRK have incomplete polybasic clusters (Fig. 3B), indicating that the NRK CNH domain underwent molecular evolution and changed its properties to a phospholipid-binding domain during the ancestor of the placental mammals. Our structural modeling showed that mNRK and cNRK CNH domains form a β-propeller structure similar to the prototypical CNH domain of yeast Rom2 (30). The mNRK CNH domain has evolved to acquire complete polybasic clusters and a larger positively charged surface in contact with the lipid bilayer. To our knowledge, the PROPPIN family is the only phospholipid-binding domain characterized by a β-propeller structure with a unique ability to bind phosphatidylinositol phosphates (PIPs) such as PI(3)P and PI(3,5)P_2_ (40). PROPPINs are endomembrane-localizing scaffold proteins for assembling protein complexes, such as autophagy machinery components (41). PROPPINs initially target membranes through nonspecific electrostatic interactions, followed by retention via PIP binding to two specific sites and membrane insertion of loop 6CD (42). In contrast, the mNRK CNH domain preferentially binds to phosphatidylserine and interacts with the PM utilizing a broader surface interface involving complete polybasic clusters. These findings highlight distinctive mechanisms of membrane recognition and interaction among phospholipid-binding domains with a β-propeller structure.

### The cellular function of the NRK CNH domain

The mechanisms by which the mNRK CNH domain promotes cell death remain unclear. mNRK plays a role in suppressing AKT activation (20, 43), an anti-apoptotic factor, and increasing JNK activation, a pro-apoptotic factor (44). However, the CNH domain alone neither inhibits AKT signaling (20), increases JNK phosphorylation, nor affects the caspase cascade (supplementary figure 3A). The mNRK CNH domain localizes at the PM throughout various stages of apoptosis, with some accumulation observed at membrane blebbing sites. The mNRK CNH domain interacts with PS (20), implying a potential role of the CNH domain not in regulating death signal but in more downstream events at the PM, for example, PS exposure.

We speculated that the mNRK CNH domain might function as a scaffold protein like PROPPINs, and the cell death-promoting activity may be mediated by some mNRK CNH domain-interacting proteins. Thus, we identified the interacting proteins, including syndecans (SDCs). SDCs can modulate survival signaling partly by associating with other PM proteins, including growth factor receptors and integrins, and several cytoplasmic signaling proteins (45, 46). SDC1 promotes and inhibits cell death in a cell type-dependent manner (47–49). SDC1 gene expression is the highest of the placental proteoglycans and is downregulated in pre-eclampsia and fetal growth restriction (50). Although detailed mechanisms of how SDCs regulate cell death remain unclear, SDCs are likely involved in the cell death machinery promoted by the mNRK CNH domain. The cell death-promoting activity of mNRK seems closely linked to removing the N-terminal sequence. The CNH domain of MINK1 interacts with its kinase domain (51), and both MINK1 and TNIK are capable of homodimerization through their central regions (52, 53), resulting in functional changes in these proteins. Functional changes triggered by protein cleavage and structural rearrangement may be a common regulatory mechanism within the GCK IV family. Therefore, cleaving off the N-terminal fragment of mNRK may disrupt intramolecular interactions or homodimer formation, thus unmasking the CNH domain to serve as a novel interaction site with cell death machinery protein(s).

### Physiological roles of the NRK CNH domain in placental development

During the later stages of pregnancy, placental weight reaches the plateau and remains constant until delivery due to decreased cell proliferation (54) and increased apoptosis (16, 55) in the spongiotrophoblast layer. Mouse NRK is highly expressed in the spongiotrophoblast layer during late pregnancy (19). We previously demonstrated that mNRK inhibits cell proliferation by modulating the CK2-PTEN-AKT pathway (20). This study found that the mNRK CNH domain promotes cell death in trophoblasts. NRK appears to regulate both cell proliferation and cell death in the spongiotrophoblast layer, contributing to the placental tissue remodeling at the late stage of pregnancy.

In conclusion, our study suggests that the NRK CNH domain has evolved to acquire polybasic clusters, enabling it to bind to phospholipids, facilitating its localization to the PM, and promoting trophoblast cell death. It is needed to elucidate detailed mechanisms underlying its cell death-promoting activity.

### Experimental procedures Plasmids

The N-terminal FLAG-tagged full-length (FL) mouse NRK (mNRK), as well as mNRK fragments spanning aa 1092-1455, mNRK ΔCNH (aa 1-1137), mNRK CNH (aa 1133-1455), and various mNRK CNH mutants were produced by utilizing the pFLAG-CMV2 (Sigma Aldrich), pcDNA5/FRT/TO (Invitrogen), and pCDH-EF1-MCS-T2A (System Biosciences) vectors. GFP-tagged versions of mNRK CNH FL, mNRK CNH A (aa 1133-1331), and mNRK CNH B (aa 1332-1455) were expressed using the pEGFP-N1 (Clontech).

### Cell culture and transfection

HEK293T and HeLa cells were cultured in DMEM, BeWO cells in Ham’s F-12, and Rcho-1 cells in RPMI. Each culture medium was supplemented with 10% FBS and 1% penicillin streptomycin. DNA transfection was conducted using polyethylenimine (PEI, Polyscience) and Lipofectamine® 3000 (Invitrogen). Apoptosis was induced by treating the cells with TNF-α (PeproTech) and cycloheximide (CHX; Nacalai Tesque).

### Lentivirus production

To generate lentivirus, HEK293T cells were transfected with pCDH-EF1-MCS-FLAG-mNRK CNH-T2A or pCDH-EF1-MCS-FLAG-mNRK CNH-18A-T2A along with psPAX2 (Addgene #12260) and pCMV-VSV-G (Addgene #8454). After a two-day transfection period, the lentivirus-containing medium was collected and filtered. HeLa cells were then exposed to this virus-containing medium in the presence of 8 μg/ml polybrene (Nacalai Tesque). Starting from 2 days post-infection, cells were cultured with 0.8 μg/ml puromycin (Invitrogen). Surviving cells were subsequently employed for experimental purposes.

### Cell survival assay (WST-8 assay)

HeLa cells (5 × 10^3^/well) were cultured for 48 h in 96-well plates, followed by a 6-hour treatment with or without TNF-α and CHX. Following treatment, 10 µL of cell count reagent SF (Nacalai Tesque) or cell counting kit-8 (Dojindo) was added, and the cells were further incubated for an additional 2 hours. The absorbance was subsequently determined at 450 nm using a microplate reader (Varioskan^TM^ LUX, Thermo Fisher Scientific).

### Cell death assay (SYTOX Green staining)

HeLa cells (5 × 10^3^/well) were cultured for 48 h in 96-well plates, followed by a 6-hour treatment with or without TNF-α and CHX. Following treatment, cells were stained with 1 μM SYTOX Green (Invitrogen) for 2 h. SYTOX Green fluorescence was measured using a fluorescence microplate reader (Varioskan^TM^ LUX, Thermo Fisher Scientific), using 485 nm excitation and 520 nm emission filters. The percentage of cell death was determined as (induced fluorescence - background fluorescence) / (maximal fluorescence - background fluorescence) × 100, with maximal fluorescence was obtained by fully permeabilizing cells using 0.1% Triton X-100.

### Immunoblotting

Cells were lysed with ice-cold lysis buffer consisting of 20 mM Tris-HCl (pH 7.4), 100 mM NaCl, 50 mM NaF, 0.5% Nonidet P-40, 1 mM EDTA, 1 mM phenylmethylsulfonyl fluoride, and protease inhibitors (1 µg/mL aprotinin, 1 µg/mL leupeptin, and 1 µg/mL pepstatin A). After centrifugation, the resulting supernatants were utilized for immunoblotting using primary antibody (anti-FLAG antibody (clone M2, Sigma-Aldrich), anti-α-Tubulin antibody (#013-25033, Wako), anti-PARP1 antibody (clone F-2, Santa Cruz), anti-caspase-3 antibody (clone 31A1067, Santa Cruz), and anti-pJNK (clone 6254, Santa Cruz) and peroxidase-conjugated anti-mouse secondary antibody (GE Healthcare). The detection of immunoblotting signals was achieved using ECL Prime Western Blotting Detection Reagents (GE Healthcare) and the ImageQuant LAS 4000 mini-imager (GE Healthcare). Densitometric analyses were carried out using the ImageJ program (version 1.52).

### Immunostaining

HeLa cells cultured on glass coverslips were transfected with PEI for 48 h and then treated with TNF-α and CHX for 1, 2, and 4 h. After treatment, cells were fixed with 4% paraformaldehyde (PFA) in PBS, permeabilized with 0.2% Triton X-100 in PBS, and blocked with 5% FBS in PBS. Cells were then incubated with a mouse anti-FLAG antibody (clone M2, Sigma-Aldrich) for 2 to 16 h at 4°C, followed by a 1-hour incubation with an Alexa Fluor 488-conjugated anti-mouse IgG secondary antibody (Invitrogen). Nuclear staining was achieved with DAPI (Nacalai Tesque) before mounting using Fluoroshield (ImmunoBioScience). Fluorescence images were captured using a laser-scanning confocal microscope (LSM 780, Carl Zeiss, Oberkochen, Germany).

For Annexin V staining, cells were cultured in a medium containing 0.25 μg/mL Annexin V (Biotium) and 2.5 mM CaCl_2_. Subsequently, fixation and immunostaining were carried out following standard protocols in the presence of a 2.5 mM CaCl_2_ in all buffers.

### Phosphatidylserine binding assay

FLAG-tagged CNH domains, encompassing wild-type and mutant variants, were produced in HEK293T cells. The cell lysates were subjected to immunoprecipitation (IP) using agarose beads coupled with anti-FLAG antibody (Sigma-Aldrich). The IP precipitates were eluted using 150 ng/mL FLAG peptides (Sigma-Aldrich) in a lysis buffer. The purified proteins were then incubated with beads coated with PS or control beads (Echelon) in a binding buffer containing 10 mM HEPES, pH 7.4, 150 mM NaCl, 0.25% Igepal at 4°C for 3 h. Afterward, the beads underwent five washes with the wash/binding buffer and were eluted with SDS-PAGE sample buffer, followed by boiling for 10 minutes. The resulting samples were subjected to SDS-PAGE and immunoblotting using anti-FLAG antibody.

### Sequence alignment

The NRK CNH amino acid sequences from multiple species using previous data (20) were aligned using ClustalW (https://clustalw.ddbj.nig.ac.jp/). In the alignment, the color blue is used to indicate the presence of polybasic or positively charged amino acids, specifically lysine (Lys or K) and arginine (Arg or R).

### *In silico* structural modeling

Since no CNH domain in the Protein Data Bank was found having sufficient sequence similarity with mNRK and cNRK CNH, we attempted *ab initio* protein structure prediction by AlphaFold2 (AF2) (56). The sequence of mNRK CNH (aa 1132–1453) was used as the input of the Colab Fold server (https://colab.research.google.com/github/sokrypton/ColabFold/). The standard parameters were used for the structure prediction and the rank1 model was taken in this study (supplementary figure 4A). Similarly, the structure model of cNRK CNH was also constructed using the sequence aa 938–1256 (supplementary figure 4B).

The predicted model of mNRK CNH had a couple of segments with low prediction accuracy, probably because of the insufficient data on the sequence-structure relationship of the CNH domain. Then, we attempted to refine the AF2 model using (1) the reference from the secondary structure prediction server JPRED (https://www.compbio.dundee.ac.uk/jpred/), (2) the knowledge of the CNH domain structural preference to β propeller (30), and (3) the modeled cNRK CNH (because this model undergoes complete β propeller structure and is in accord with the secondary structure prediction by JPRED). Firstly, the lack of β sheets in aa 1333– 1372 (red in supplementary figure 4A) was changed to three β sheets by homology modeling to the AF2 model of cNRK CNH (orange in supplementary figure 4B). The AF2 model also has a collapsed region, probably because of insufficient structure data with sequence similarity with mNRK CNH. The C-terminal helix was also remodeled as a loop, and the topology of the Exon25 structure was adjusted to align with the AF2 model of bison NRK CNH.

Then, we performed the MD simulation of the model in solution. A rectangular simulation box was constructed with a margin of 12 Å to the boundary of the simulation box, resulting in a dimension of 82 Å × 95 Å × 89 Å. The solution system contained 16735 TIP3P water molecules (57) together with 30 sodium ions and 40 chloride ions corresponding to ∼0.15 nM ion concentration, resulting in 56305 atoms in total. AMBER ff14SB was applied for the potential energy of the all-atom protein (58, 59). The MD simulations were performed by AMBER 20 (60) under constant temperature and pressure (NPT) conditions at T = 300 K and P = 1 atm using Berendsen’s thermostat and barostat (61) at a relaxation time of 1 ps and using the particle mesh Ewald method (62) for the electrostatic interactions. The simulation length was 1 μs and a 2-fs time step using constraining bonds involving hydrogen atoms via the SHAKE algorithm (63). The simulation of the cNRK CNH model was also carried out as described above (the solution system contained 14026 TIP3P waters, 30 sodium ions, 38 chloride ions, and the overall 47295 atoms in a rectangular box of 79 Å × 82 Å × 90 Å.).

To construct the CNH domain interacting with the lipid bilayer, we predicted the rotational and translational position of the peripheral protein in membranes using the PPM 3.0 Web Server (https://opm.phar.umich.edu/ppm_server) (64). The Membrane Builder module of CHARMM-GUI (https://www.charmm-gui.org/) was then applied to construct the full model containing palmitoyl-oleoyl phosphatidylserine (POPS) lipid bilayer, ions, and waters (65). The mNRK CNH system contained 300 POPSs, 27,084 TIP3P waters, 362 potassium ions, 72 chloride ions (that are equal to 0.15 nM ion concentration), and the overall 125,816 atoms in a rectangular box of 98 Å × 98 Å × 139 Å. The cNRK CNH system contained 270 POPSs, 23,656 TIP3P waters, 325 potassium ions, 63 chloride ions, and overall 110,825 atoms in a rectangular box of 92 Å × 92 Å × 137 Å. MD simulations were conducted with AMBER 20 using the generated input files. Three runs were executed for each system, and the trajectories of the 100-ns product runs were used for analysis.

### LC-MS/MS analysis

Parental HeLa cells and HeLa cells stably expressing the FLAG-CNH domain were cultured for 48 h. The cells were treated with or without TNF-α and CHX and incubated for 2 h. The lysates were subjected to immunoprecipitation using anti-FLAG antibody-conjugated agarose beads (Sigma-Aldrich). The precipitates were eluted with 150 ng/mL FLAG peptides (Sigma-Aldrich) in lysis buffer and subjected to reduction and alkylation processes. The eluted proteins were then reduced with TCEP (tris (2-carboxyethyl) phosphine) (Thermo Fisher Scientific) at a final concentration of 25 mM Tris for 15 min at 20–25 °C and alkylated with 55 mM iodoacetamide for 45 min at 20–25 °C in a light-shielded condition. Next, the proteins were concentrated by acetone precipitation for at least 1 h. The proteins were then subjected to SDS-PAGE. After protein separation by SDS-PAGE, silver staining was performed according to the protocol provided by WAKO. The gels were cut (∼1 cm) into pieces followed by de-staining process and proceed to in-gel digestion protocol. Then, the gels were dehydrated by 100 mM NH_4_HCO_3_ solution by 50% CH_3_CN in 100 mM NH_4_HCO_3_ solution. After the solution was removed, trypsin (MS grade; Thermo Fisher Scientific) solution was added to each tube and incubated overnight at 37 °C. The resulting solution was quenched with aqueous Trifluoroacetic acid (TFA) solution (final concentration: 0.1%) and desalted using Sep-Pak C18 96-well plates (Waters, Manchester, UK) and dried under reduced pressure. The sample was suspended in 0.1% TFA and 2% CH_3_CN, and then subjected to liquid chromatography-tandem mass spectrometry (LC-MS/MS) analysis using a nano-liquid chromatography column (EASY-nLC 1000; Thermo Fisher Scientific) coupled to the Q Exactive mass spectrometer (Thermo Fisher Scientific). Data analysis was performed using the Proteome Discoverer 2.4 software embedded with the Sequest algorithm (Thermo Fisher Scientific).

## Data availability

Data supporting the study of molecular dynamic simulations are available from the corresponding authors upon reasonable request. Other data are included within this article.

## Author contributions

B.L. and T.F. conceptualization; B.L., T.F., K.M., and A.K. methodology; B.L., K.S., T.F., K.M., and A.K. investigation; Y.S., Y.H., and E.M. resources; B.L., K.M., and T.F. writing–original draft; B.L., K.M., A.K., and T.F. writing-review and editing; K.M., M.K., M.S.H.W., and T.F. funding acquisition.

## Funding and additional information

We thank the Biomaterials Analysis Division, Tokyo Institute of Technology, for DNA sequencing and confocal microscopy analysis and Dr. Tatsuya Niwa for helping our LC-MS/MS analysis. We also thank Dr. Satoshi Tanaka (The University of Tokyo) for kindly providing Rcho-1 cell lines. This research received financial support from the Post-Doctoral Program of the Directorate of Research, Universitas Gadjah Mada, and the team from the Office of Quality Assurance, Universitas Gadjah Mada under Grant No. 3662/UN1.P.II/Dit-Lit/PT.01.03/2023 (to B.L. and M.S.H.W). Additionally, funding is extended through JSPS KAKENHI under Grant No. JP22K06039 (to T.F.) and the Research Support Project for Life Science and Drug Discovery (BINDS) from AMED under Grant No. JP23ama121023 (to K.M.).

---

**Supplementary figure 1.**
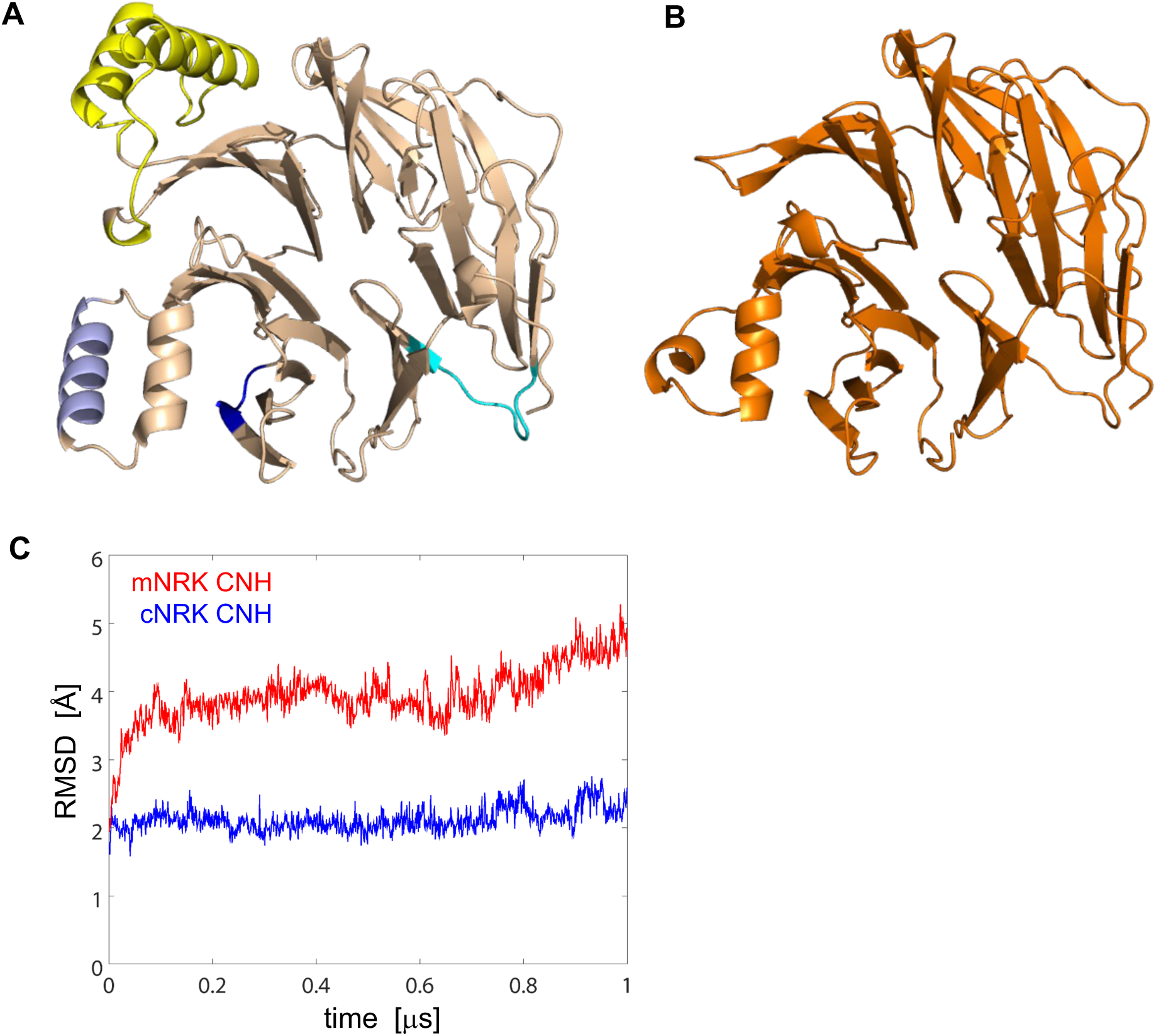
Structural modeling of the mNRK and cNRK CNH domains. *A, B,* Models of the mNRK (A) and cNRK (B) CNH domains were constructed as described in the “Experimental procedure.” In A, sites 1-3 of the polybasic clusters are colored in blue, light blue, and cyan, respectively, and the region encoded by the exon 25 is shown in yellow. *C,* The overall Cα-RMSD values during the 1-μs MD simulations are plotted for mNRK (red) and cNRK (blue) CNH domains. The overall Cα-RMSD < 5 Å from the initial models indicates substantial stability. The larger Cα-RMSD for the mNRK CNH domain comes from the flexibility of the region encoded in the exon 25.

**Supplementary figure 2.**
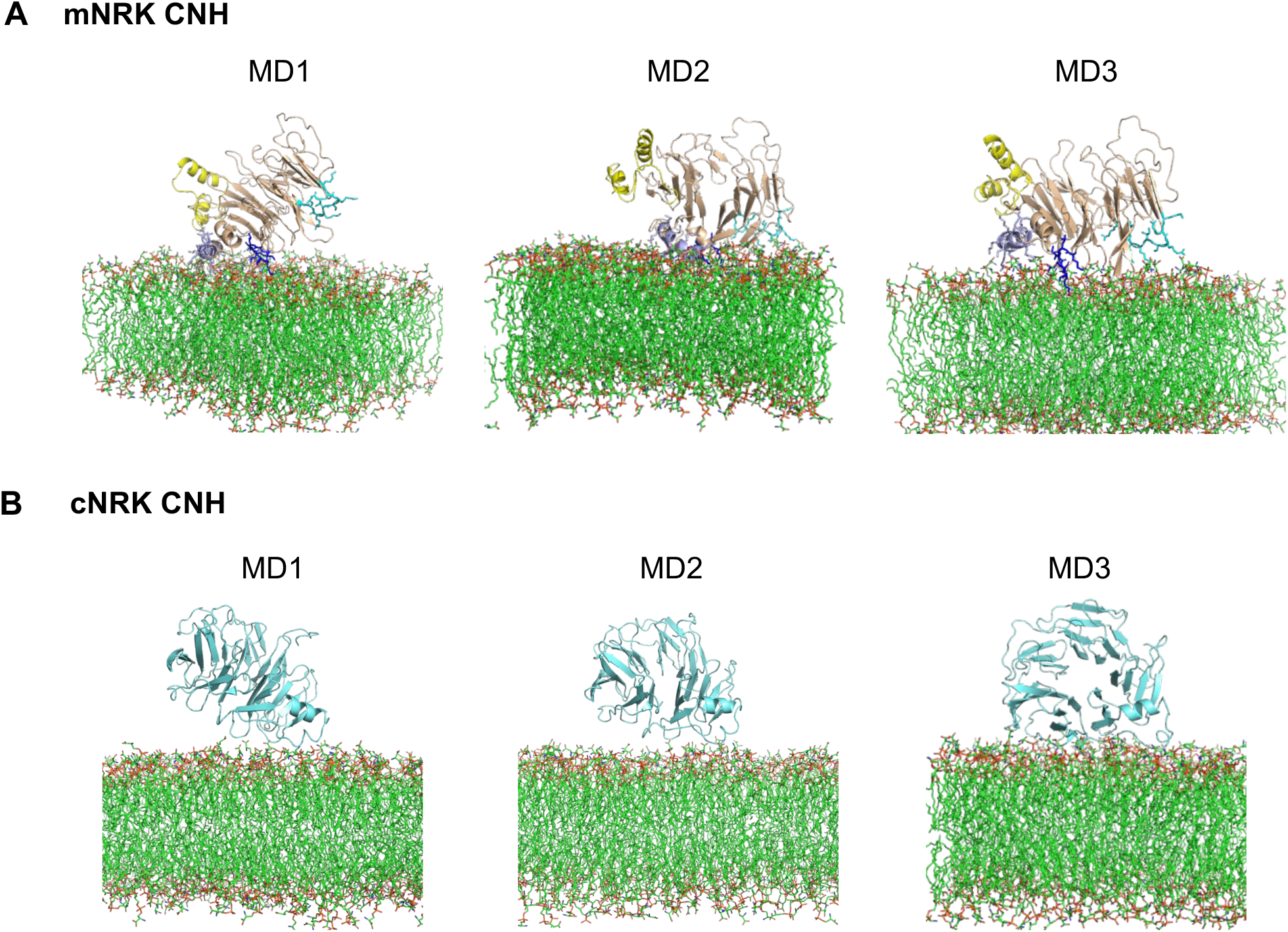
Representative structures of the membrane-bound mNRK CNH (upper) and cNRK CNH (lower) domains taken from the three MD simulations.

**Supplementary figure 3.**
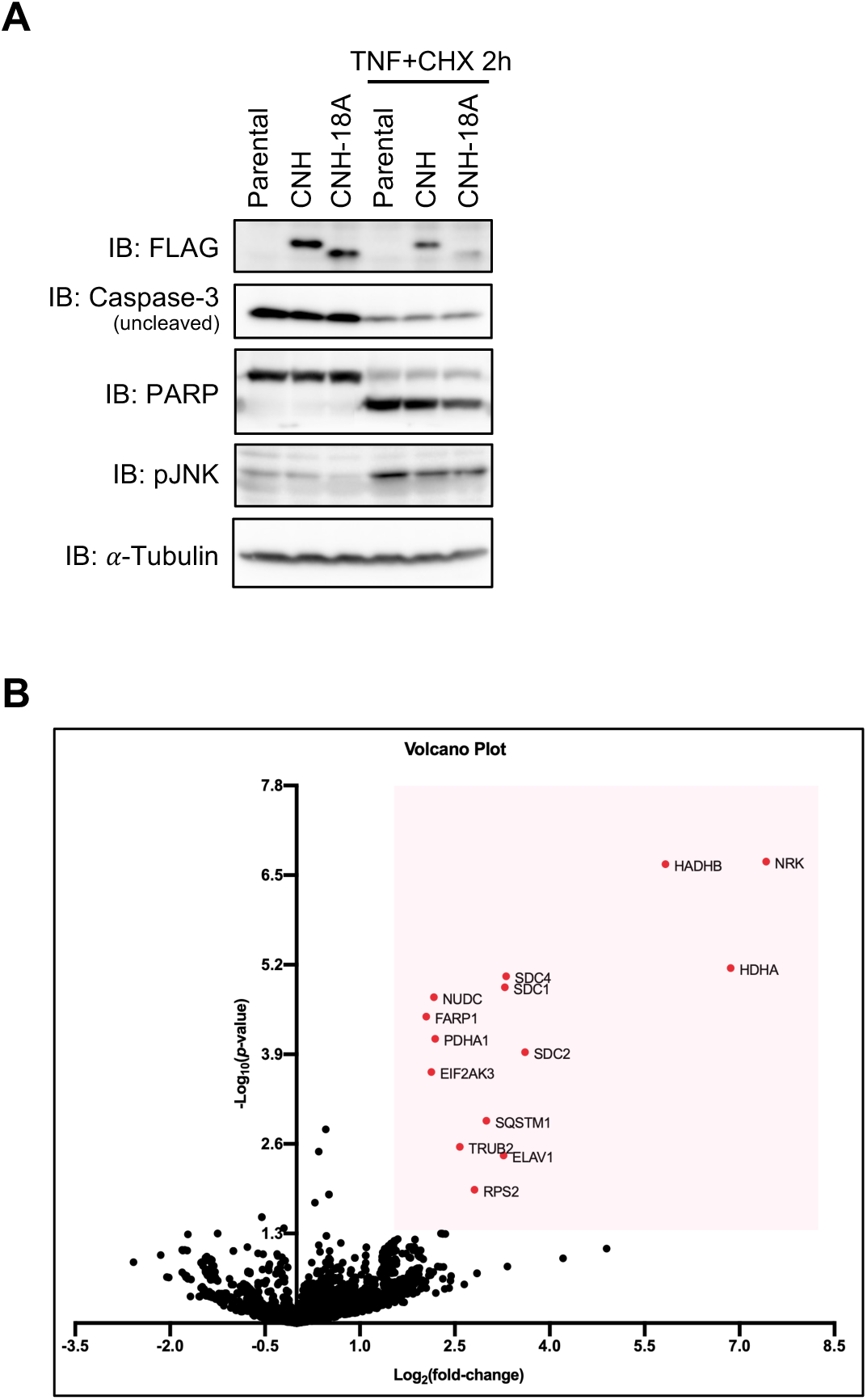
Effects of the mNRK CNH domain on caspase-3 and PARP cleavage and the mNRK CNH domain-interacting proteins during apoptosis. *A,* HeLa cells stably expressing the mNRK CNH domain and CNH-18A were treated with TNF-*α* (10 ng/mL) and CHX (10 *μ*g/mL) for 2 h and subjected to immunoblotting using indicated antibodies. *B,* HeLa cells stably expressing FLAG-tagged CNH domain and parental cells (control) were treated with TNF-*α* (10 ng/mL) and CHX (10 *μ* g/mL) for 2 h. Cell lysates were subjected to immunoprecipitation with anti-FLAG antibody, followed by SDS-PAGE and silver staining. Co-immunoprecipitated proteins were treated with in-gel digestion and identified by LC-MS/MS analysis. The volcano plot shows the result of the MS analysis; the vertical axis indicates each protein’s relative intensity (fold of control, log_2_ transformed), and the horizontal axis indicates the *p*-value (-log_10_ transformed). The red dots indicate significant proteins.

**Supplementary figure 4.**
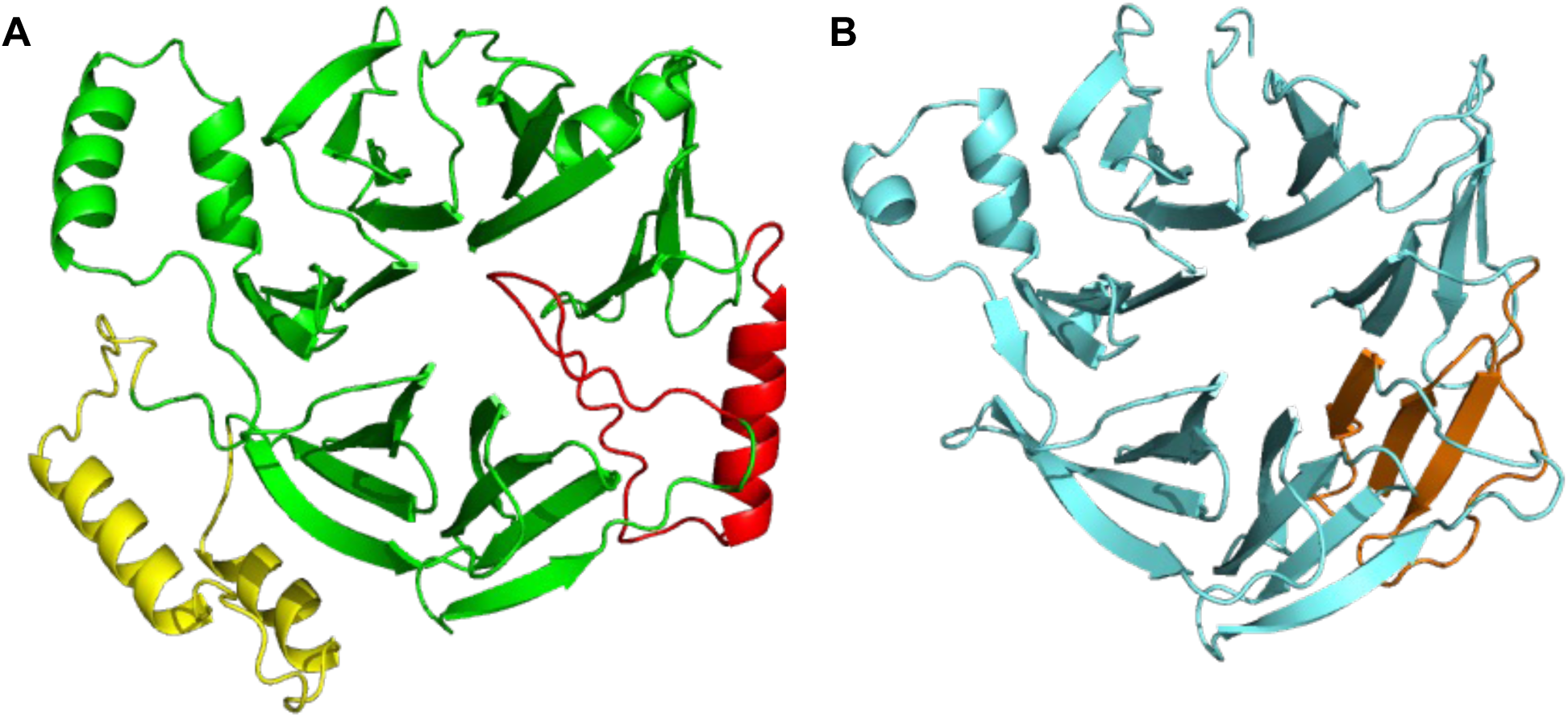
AF2 structures of the mNRK and cNRK CNH domains. *A, B,* Models of the mNRK (A) and cNRK (B) CNH domains were constructed using AlphaFold2 (AF2) (56). The predicted region that lacks the β sheets for mNRK (aa 250-289) is shown by red, and the corresponding region for cNRK is shown by orange. Exon25 is shown by yellow.

**Supplementary table 1.** The list of mNRK CNH domain-interacting proteins during apoptosis.

| Gene | Protein name | Description |
| --- | --- | --- |
| HADHA, HADHB | Hydroxyl Coenzyme A (CoA) Dehydrogenase subunits $\alpha$ and $\beta$ | Mitochondrial trifunctional proteins which catalyzes the last three steps of fatty acid oxidation |
| SDC1,2,4 | Syndecan isoform 1, 2, and 4 | Transmembrane proteins which play roles in cell adhesion, migration, and signaling |
| NUDC | Nuclear Distribution C, Dynein Complex Regulator | Nuclear protein which involves in mitosis and cytokinesis |
| FARP1 | FERM, ARH/RhoGEF And Pleckstrin Domain Protein 1 | Cytoskeleton-associated proteins at plasma membrane which can activate Rho GTPases |
| PDHA1 | Pyruvate Dehydrogenase E1 Subunit Alpha 1 | Nuclear-encoded mitochondrial multienzyme complex that catalyzes the conversion of pyruvate to acetyl-CoA and CO |
| EIF2AK3 | Eukaryotic Translation Initiation Factor 2 Alpha Kinase 3 | Endoplasmic reticulum (ER)-resident protein that mediates signal transduction during ER stress |
| SQSTM1 | Sequestosome 1/p62 | Nucleocytoplasmic adapter protein involved in the transportation, degradation and destruction of various proteins |
| TRUB2 | TruB Pseudouridine Synthase Family Member 2 | Protein acts as upstream of or within RNA processing and pseudouridine synthesis |
| ELAV1 | ELAV Like RNA Binding Protein 1 | RNA-binding protein that binds to the 3'-UTR region of mRNAs and increases their stability |
| RPS2 | Ribosomal Protein S2 | Component of the ribosome, a large ribonucleoprotein complex responsible for the synthesis of proteins in the cell |

## References

1. Rossant, J., and Cross, J. C. (2001) Placental development: Lessons from mouse mutants. Nat Rev Genet. 2, 538–548

2. John, R., and Hemberger, M. (2012) A placenta for life. Reproductive BioMedicine Online. 25, 5–11

3. Moffett, A., and Loke, C. (2006) Immunology of placentation in eutherian mammals. Nat Rev Immunol. 6, 584–594

4. Huppertz, B., Frank, H. G., Kingdom, J. C., Reister, F., and Kaufmann, P. (1998) Villous cytotrophoblast regulation of the syncytial apoptotic cascade in the human placenta. Histochem Cell Biol. 110, 495–508

5. Gong, J.-S., and Kim, G. J. (2014) The role of autophagy in the placenta as a regulator of cell death. Clin Exp Reprod Med. 41, 97–107

6. Mayhew, T. M., Leach, L., McGee, R., Wan Ismail, W., Myklebust, R., and Lammiman, M. J. (1999) Proliferation, Differentiation and Apoptosis in Villous Trophoblast at 13–41 Weeks of Gestation (Including Observations on Annulate Lamellae and Nuclear Pore Complexes). Placenta. 20, 407–422

7. Dumolt, J. H., Powell, T. L., and Jansson, T. (2021) Placental function and the development of fetal overgrowth and fetal growth restriction. Obstetrics and gynecology clinics of North America. 48, 247

8. Lee, E. D., and Mistry, H. D. (2022) Placental Related Disorders of Pregnancy. International Journal of Molecular Sciences. 23, 3519

9. Fuchs, Y., and Steller, H. (2011) Programmed cell death in animal development and disease. Cell. 147, 742–758

10. Vaux, D. L., and Korsmeyer, S. J. (1999) Cell Death in Development. Cell. 96, 245–254

11. Kerr, J. F., Wyllie, A. H., and Currie, A. R. (1972) Apoptosis: a basic biological phenomenon with wide-ranging implications in tissue kinetics. Br J Cancer. 26, 239–257

12. Sharp, A. N., Heazell, A. E. P., Crocker, I. P., and Mor, G. (2010) Placental Apoptosis in Health and Disease. Am J Reprod Immunol. 64, 159–169

13. Athapathu, H., Jayawardana, M. a. J., and Senanayaka, L. (2003) A study of the incidence of apoptosis in the human placental cells in the last weeks of pregnancy. J Obstet Gynaecol. 23, 515–517

14. Smith, S. C., and Baker, P. N. (1999) Placental apoptosis is increased in post-term pregnancies. Br J Obstet Gynaecol. 106, 861–862

15. Detmar, J., Rovic, I., Ray, J., Caniggia, I., and Jurisicova, A. (2019) Placental cell death patterns exhibit differences throughout gestation in two strains of laboratory mice. Cell Tissue Res. 378, 341–358

16. Waddell, B. J., Hisheh, S., Dharmarajan, A. M., and Burton, P. J. (2000) Apoptosis in Rat Placenta Is Zone-Dependent and Stimulated by Glucocorticoids1. Biology of Reproduction. 63, 1913–1917

17. Kanai-Azuma, M., Kanai, Y., Okamoto, M., Hayashi, Y., Yonekawa, H., and Yazaki, K. (1999) Nrk: a murine X-linked NIK (Nck-interacting kinase)-related kinase gene expressed in skeletal muscle. Mechanisms of Development. 89, 155–159

18. Liu, G., Zhang, C., Wang, Y., Dai, G., Liu, S.-Q., Wang, W., Pan, Y.-H., Ding, J., and Li, H. (2021) New exon and accelerated evolution of placental gene Nrk occurred in the ancestral lineage of placental mammals. Placenta. 114, 14–21

19. Denda, K., Nakao-Wakabayashi, K., Okamoto, N., Kitamura, N., Ryu, J.-Y., Tagawa, Y., Ichisaka, T., Yamanaka, S., and Komada, M. (2011) Nrk, an X-linked Protein Kinase in the Germinal Center Kinase Family, Is Required for Placental Development and Fetoplacental Induction of Labor. J Biol Chem. 286, 28802–28810

20. Lestari, B., Naito, S., Endo, A., Nishihara, H., Kato, A., Watanabe, E., Denda, K., Komada, M., and Fukushima, T. (2022) Placental Mammals Acquired Functional Sequences in NRK for Regulating the CK2–PTEN–AKT Pathway and Placental Cell Proliferation. Molecular Biology and Evolution. 39, msab371

21. Vento-Tormo, R., Efremova, M., Botting, R. A., Turco, M. Y., Vento-Tormo, M., Meyer, K. B., Park, J.-E., Stephenson, E., Polański, K., Goncalves, A., Gardner, L., Holmqvist, S., Henriksson, J., Zou, A., Sharkey, A. M., Millar, B., Innes, B., Wood, L., Wilbrey-Clark, A., Payne, R. P., Ivarsson, M. A., Lisgo, S., Filby, A., Rowitch, D. H., Bulmer, J. N., Wright, G. J., Stubbington, M. J. T., Haniffa, M., Moffett, A., and Teichmann, S. A. (2018) Single-cell reconstruction of the early maternal-fetal interface in humans. Nature. 563, 347–353

22. Cross, J. C., Nakano, H., Natale, D. R. C., Simmons, D. G., and Watson, E. D. (2006) Branching morphogenesis during development of placental villi. Differentiation. 74, 393–401

23. Carter, A. M. (2007) Animal models of human placentation--a review. Placenta. 28 Suppl A, S41–47

24. Yomogita, H., Ito, H., Hashimoto, K., Kudo, A., Fukushima, T., Endo, T., Hirate, Y., Akimoto, Y., Komada, M., Kanai, Y., Miyasaka, N., and Kanai-Azuma, M. (2022) A possible function of *Nik-related kinase* in the labyrinth layer of delayed delivery mouse placentas. Journal of Reproduction and Development. advpub, 2022–120

25. Dan, I., Watanabe, N. M., and Kusumi, A. (2001) The Ste20 group kinases as regulators of MAP kinase cascades. Trends in Cell Biology. 11, 220–230

26. Delpire, E. (2009) The mammalian family of sterile 20p-like protein kinases. Pflugers Arch - Eur J Physiol. 458, 953–967

27. Kakinuma, H., Inomata, H., and Kitamura, N. (2005) Enhanced JNK activation by NESK without kinase activity upon caspase-mediated cleavage during apoptosis. Cellular Signalling. 17, 1439–1448

28. Nagata, S., Suzuki, J., Segawa, K., and Fujii, T. (2016) Exposure of phosphatidylserine on the cell surface. Cell Death Differ. 23, 952–961

29. Rogers, C., Fernandes-Alnemri, T., Mayes, L., Alnemri, D., Cingolani, G., and Alnemri, E. S. (2017) Cleavage of DFNA5 by caspase-3 during apoptosis mediates progression to secondary necrotic/pyroptotic cell death. Nat Commun. 8, 14128

30. Bartual, S. G., Wei, W., Zhou, Y., Pravata, V. M., Fang, W., Yan, K., Ferenbach, A. T., Lockhart, D. E. A., and van Aalten, D. M. F. (2021) The citron homology domain as a scaffold for Rho1 signaling. Proceedings of the National Academy of Sciences. 118, e2110298118

31. Msheik, H., El Hayek, S., Bari, M. F., Azar, J., Abou-Kheir, W., Kobeissy, F., Vatish, M., and Daoud, G. (2019) Transcriptomic profiling of trophoblast fusion using BeWo and JEG-3 cell lines. Molecular Human Reproduction. 25, 811–824

32. Koh, Y. Q., Chan, H.-W., Nitert, M. D., Vaswani, K., Mitchell, M. D., and Rice, G. E. (2014) Differential response to lipopolysaccharide by JEG-3 and BeWo human choriocarcinoma cell lines. European Journal of Obstetrics & Gynecology and Reproductive Biology. 175, 129–133

33. Soares, M. J., Chakraborty, D., Karim Rumi, M. A., Konno, T., and Renaud, S. J. (2012) Rat placentation: an experimental model for investigating the hemochorial maternal-fetal interface. Placenta. 33, 233–243

34. Iwatsuki, K., Shinozaki, M., Sun, W., Yagi, S., Tanaka, S., and Shiota, K. (2000) A Novel Secretory Protein Produced by Rat Spongiotrophoblast1. Biology of Reproduction. 62, 1352–1359

35. Taira, K., Umikawa, M., Takei, K., Myagmar, B.-E., Shinzato, M., Machida, N., Uezato, H., Nonaka, S., and Kariya, K. (2004) The Traf2- and Nck-interacting kinase as a putative effector of Rap2 to regulate actin cytoskeleton. J Biol Chem. 279, 49488–49496

36. Machida, N., Umikawa, M., Takei, K., Sakima, N., Myagmar, B.-E., Taira, K., Uezato, H., Ogawa, Y., and Kariya, K.-I. (2004) Mitogen-activated protein kinase kinase kinase kinase 4 as a putative effector of Rap2 to activate the c-Jun N-terminal kinase. J Biol Chem. 279, 15711–15714

37. Nonaka, H., Takei, K., Umikawa, M., Oshiro, M., Kuninaka, K., Bayarjargal, M., Asato, T., Yamashiro, Y., Uechi, Y., Endo, S., Suzuki, T., and Kariya, K.-I. (2008) MINK is a Rap2 effector for phosphorylation of the postsynaptic scaffold protein TANC1. Biochem Biophys Res Commun. 377, 573–578

38. Kawabe, H., Neeb, A., Dimova, K., Young, S. M., Takeda, M., Katsurabayashi, S., Mitkovski, M., Malakhova, O. A., Zhang, D.-E., Umikawa, M., Kariya, K., Goebbels, S., Nave, K.-A., Rosenmund, C., Jahn, O., Rhee, J., and Brose, N. (2010) Regulation of Rap2A by the Ubiquitin Ligase Nedd4-1 Controls Neurite Development. Neuron. 65, 358–372

39. Duncan, E. D., Han, K.-J., Trout, M. A., and Prekeris, R. (2022) Ubiquitylation by Rab40b/Cul5 regulates Rap2 localization and activity during cell migration. J Cell Biol. 221, e202107114

40. Baskaran, S., Ragusa, M. J., Boura, E., and Hurley, J. H. (2012) Two-site recognition of phosphatidylinositol 3-phosphate by PROPPINs in autophagy. Mol Cell. 47, 339–348

41. Marquardt, L., and Thumm, M. (2023) Autophagic and non-autophagic functions of the Saccharomyces cerevisiae PROPPINs Atg18, Atg21 and Hsv2. Biological Chemistry. 404, 813–819

42. Busse, R. A., Scacioc, A., Krick, R., Pérez-Lara, Á., Thumm, M., and Kühnel, K. (2015) Characterization of PROPPIN-Phosphoinositide Binding and Role of Loop 6CD in PROPPIN-Membrane Binding. Biophys J. 108, 2223–2234

43. Naito, S., Fukushima, T., Endo, A., Denda, K., and Komada, M. (2020) Nik-related kinase is targeted for proteasomal degradation by the chaperone-dependent ubiquitin ligase CHIP. FEBS Lett. 594, 1778–1786

44. Nakano, K., Yamauchi, J., Nakagawa, K., Itoh, H., and Kitamura, N. (2000) NESK, a member of the germinal center kinase family that activates the c-Jun N-terminal kinase pathway and is expressed during the late stages of embryogenesis. J Biol Chem. 275, 20533–20539

45. Afratis, N. A., Nikitovic, D., Multhaupt, H. A. B., Theocharis, A. D., Couchman, J. R., and Karamanos, N. K. (2017) Syndecans – key regulators of cell signaling and biological functions. The FEBS Journal. 284, 27–41

46. Khotskaya, Y. B., Dai, Y., Ritchie, J. P., MacLeod, V., Yang, Y., Zinn, K., and Sanderson, R. D. (2009) Syndecan-1 is required for robust growth, vascularization, and metastasis of myeloma tumors in vivo. J Biol Chem. 284, 26085–26095

47. Boeddeker, S. J., Baston-Buest, D. M., Altergot-Ahmad, O., Kruessel, J. S., and Hess, A. P. (2014) Syndecan-1 knockdown in endometrial epithelial cells alters their apoptotic protein profile and enhances the inducibility of apoptosis. Molecular Human Reproduction. 20, 567–578

48. Boeddeker, S. J., Baston-Buest, D. M., Fehm, T., Kruessel, J., and Hess, A. (2015) Decidualization and Syndecan-1 Knock Down Sensitize Endometrial Stromal Cells to Apoptosis Induced by Embryonic Stimuli. PLOS ONE. 10, e0121103

49. Hilgers, K., Ibrahim, S. A., Kiesel, L., Greve, B., Espinoza-Sánchez, N. A., and Götte, M. (2022) Differential Impact of Membrane-Bound and Soluble Forms of the Prognostic Marker Syndecan-1 on the Invasiveness, Migration, Apoptosis, and Proliferation of Cervical Cancer Cells. Frontiers in Oncology. [online] https://www.frontiersin.org/articles/10.3389/fonc.2022.803899 (Accessed January 14, 2023)

50. Oravecz, O., Balogh, A., Romero, R., Xu, Y., Juhasz, K., Gelencser, Z., Xu, Z., Bhatti, G., Pique-Regi, R., Peterfia, B., Hupuczi, P., Kovalszky, I., Murthi, P., Tarca, A. L., Papp, Z., Matko, J., and Than, N. G. (2022) Proteoglycans: Systems-Level Insight into Their Expression in Healthy and Diseased Placentas. International Journal of Molecular Sciences. 23, 5798

51. Lim, J., Lennard, A., Sheppard, P. W., and Kellie, S. (2003) Identification of residues which regulate activity of the STE20-related kinase hMINK. Biochemical and Biophysical Research Communications. 300, 694–698

52. Mikryukov, A., and Moss, T. (2012) Agonistic and antagonistic roles for TNIK and MINK in non-canonical and canonical Wnt signalling. PLoS One. 7, e43330

53. Kukimoto-Niino, M., Shirouzu, M., and Yamada, T. (2022) Structural Insight into TNIK Inhibition. International Journal of Molecular Sciences. 23, 13010

54. Coan, P. m., Conroy, N., Burton, G. j., and Ferguson-Smith, A. c. (2006) Origin and characteristics of glycogen cells in the developing murine placenta. Developmental Dynamics. 235, 3280–3294

55. Ishikawa, H., Seki, R., Yokonishi, S., Yamauchi, T., and Yokoyama, K. (2006) Relationship between fetal weight, placental growth and litter size in mice from mid- to late-gestation. Reproductive Toxicology. 21, 267–270

56. Jumper, J., Evans, R., Pritzel, A., Green, T., Figurnov, M., Ronneberger, O., Tunyasuvunakool, K., Bates, R., Žídek, A., Potapenko, A., Bridgland, A., Meyer, C., Kohl, S. A. A., Ballard, A. J., Cowie, A., Romera-Paredes, B., Nikolov, S., Jain, R., Adler, J., Back, T., Petersen, S., Reiman, D., Clancy, E., Zielinski, M., Steinegger, M., Pacholska, M., Berghammer, T., Bodenstein, S., Silver, D., Vinyals, O., Senior, A. W., Kavukcuoglu, K., Kohli, P., and Hassabis, D. (2021) Highly accurate protein structure prediction with AlphaFold. Nature. 596, 583–589

57. Jorgensen, W. L., Chandrasekhar, J., Madura, J. D., Impey, R. W., and Klein, M. L. (1983) Comparison of simple potential functions for simulating liquid water. The Journal of Chemical Physics. 79, 926–935

58. Cornell, W. D., Cieplak, P., Bayly, C. I., Gould, I. R., Merz, K. M., Ferguson, D. M., Spellmeyer, D. C., Fox, T., Caldwell, J. W., and Kollman, P. A. (1995) A Second Generation Force Field for the Simulation of Proteins, Nucleic Acids, and Organic Molecules. J. Am. Chem. Soc. 117, 5179–5197

59. Maier, J. A., Martinez, C., Kasavajhala, K., Wickstrom, L., Hauser, K. E., and Simmerling, C. (2015) ff14SB: Improving the Accuracy of Protein Side Chain and Backbone Parameters from ff99SB. J Chem Theory Comput. 11, 3696– 3713

60. Case, D. A., Cheatham, T. E., Darden, T., Gohlke, H., Luo, R., Merz, K. M., Onufriev, A., Simmerling, C., Wang, B., and Woods, R. J. (2005) The Amber biomolecular simulation programs. J Comput Chem. 26, 1668–1688

61. Juffer, A. H., and Berendsen, H. J. C. (1991) The electric potential of a macromolecule in a solvent: A fundamental approach. Journal of Computational Physics. 93, 251

62. Pastor, R. W., and Karplus, M. (1988) Parametrization of the friction constant for stochastic simulations of polymers. J. Phys. Chem. 92, 2636–2641

63. Ryckaert, J.-P., Ciccotti, G., and Berendsen, H. J. C. (1977) Numerical integration of the cartesian equations of motion of a system with constraints: molecular dynamics of n-alkanes. Journal of Computational Physics. 23, 327– 341

64. Lomize, M. A., Pogozheva, I. D., Joo, H., Mosberg, H. I., and Lomize, A. L. (2012) OPM database and PPM web server: resources for positioning of proteins in membranes. Nucleic Acids Research. 40, D370–D376

65. Qi, Y., Cheng, X., Lee, J., Vermaas, J. V., Pogorelov, T. V., Tajkhorshid, E., Park, S., Klauda, J. B., and Im, W. (2015) CHARMM-GUI HMMM Builder for Membrane Simulations with the Highly Mobile Membrane-Mimetic Model. Biophys J. 109, 2012–2022

